# Small RNA expression pattern in multiply inbred lines and their hybrids of maize embryo

**DOI:** 10.1101/565911

**Authors:** Yong-Xin Liu

**Affiliations:** The State Key Laboratory of Plant Genomics, Institute of Genetics and Developmental Biology, Chinese Academy of Sciences, Beijing 100101, China; University of Chinese Academy of Sciences, Beijing 100049, China

**Keywords:** maize, heterosis, sRNA-seq, small RNA cluster, retrotransposon

## Abstract

Heterosis, also known as hybrid vigor or transgression, is the phenomenon wherein an F1 hybrid produced from crossing two cultivars of the same or different species shows superior phenotypes. Heterosis in maize has been found and applied in breeding for more than one hundred years. However, its underlying molecular mechanisms are still poorly understood. To investigate whether small RNAs (sRNAs) participate in the regulation of heterosis, we profiled the sRNA expression patterns in the germ seeds of five inbred lines and theirs three F1 hybrids using high-throughput sequencing technology. The sequencing result show maize sRNAs are enriched in 22-nt length. Nearly 90% of 22-nt small RNA dominated clusters (SRCs) are in repeat regions, which mainly originated from Gypsy and Copia in retrotransposon. About 25% differentially expressed SRCs exist between parents, and hybrid gain almost all differentially expressed 22-nt SRCs. Twenty-four-nt sRNA also enriched in maize, which showed great sequence diversity and overall low expression across the genome. More than half of 24-nt SRCs originate from repeat, and 80% of them come from DNA transposon. Nearly 30% of 24-nt SRCs located in genes or their flanking regions, especially in flanking regions of “lipid metabolic process” and “macromolecule modification” related genes. Several thousands 24-nt SRCs are paternal or maternal specific expressed, and hybrids gain only half of differentially expressed 24-nt SRCs. Hundreds of 24-nt SRCs show high parent or above high parent expression pattern in different hybrids, and them mainly from Tourist, haT, and CACTA in DNA transposon and enrich nearby “tRNA aminoacylation for protein translation” related genes. Also, some 21-nt SRCs show conserved expression pattern in low parent. They were enriched in anti-sense region of some genes, which functions are about oxidative phosphorylation and translation. miRNAs have a global downregulated tendency in hybrids. zma-miR408-5p, zma-miR1432-5p and zma-miR528-5p are significant downregulated in each hybrid, and this phenomenon may cause their target genes more stable and contribute to hybrid vigor. Taken together, our results illustrated that sRNAs may contribute to heterosis at the very early stage of seed germination through repressing of retrotransposon activity, regulation gene activity at gene and genic flanking regions, and promotion some gene expressions by downregulated miRNAs.

## Introduction

Heterosis, also known as hybrid vigor or transgression, is the phenomenon wherein an F1 hybrid produced from crossing two cultivars of the same or different species shows superior phenotypes^1^. The phenomenon of heterosis, depends on genetic variation between parents and altered genetic states in their offspring^2^. Maize is a suitable model for research the genetic mechanism of heterosis, because it includes highly diversified phenotypes, allele, and the high-quality genome^3, 4^. In maize, heterosis has been found and applied in breeding for more than one hundred years^5^. Many heterosis-related QTLs have been mapped on maize genome, and many potential heterosis-related genes have been reported^6^. However, due to its so complex characters, the molecular mechanism of heterosis in maize is still poorly understood.

Small RNAs (sRNAs) are mainly 20∼24 nucleotide (nt) length RNA, which mainly functions are regulate gene expression and maintain genome integrity^7-9^. sRNA mainly included three groups, small interfering RNAs (siRNAs), microRNAs (miRNAs) and trans-acting siRNAs (ta-siRNAs) in plant^10, 11^. The 24-nt siRNAs are mainly derived from transposable elements (TEs) and repeat regions, and interact with ARGONAUTE4 (AGO4) and lead to gene silencing and/or RNA-directed DNA methylation (RdDM) at target loci^12^. Groszmann, et al. ^13^ reported 24-nt siRNA great downregulated in *Arabidopsis* hybrids suggest an epigenetic contribution to hybrid vigor. RNA directed DNA methylation (RdDM) is important in maintain steady, due to 85% of maize genome is TEs^3, 12^.

The magnitude of hybrid vigor in maize is relatively high and observed throughout its whole life cycle. Seed germination plays a pivotal role during the life cycle of plants, and many papers reported hybrid show significant different from their parents in miRNAs and proteins at this stage^6, 14^. Barber, et al. ^2^ reported repeat associated sRNAs vary among parents and following hybridization in maize of B73 and Mol17. Ding, et al. ^6^ microRNA (miRNA) prefer downregulated in hybrid of maize seed germination. These works were only used one hybrid and their parents (inbred lines), and their finding about sRNAs different between hybrid and their parents is hardly to say a conserved mechanism contributed to heterosis.

In this study, to learn more about how sRNAs change in hybridization of maize and whether exist a conserved expression pattern between hybrid and their parents, we sequenced sRNAs from the germinated seed under imbibition 24 hours from three cross combinations (include five inbred lines and three their hybrids), which are main variety in China. We identified small RNA cluster (SRCs) across maize genome, and found 22-nt dominant SRCs account for 40% and 24-nt dominant SRCs account for 48%. 22-nt SRCs mainly originated from Gypsy and Copia super families in retrotransposon, but without consistent expression trend in different hybrids. 24-nt SRCs not only enriched in DNA transposon region, but also 40% of them located in genes and their flanking (±2kb) regions, especially enriched in flanking regions of lipid metabolic process and macromolecule modification related genes, and upregulated in hybrid, which may contribute to provide more material and energy for germ seed. Maize hybrids have more SRCs with expression levels of high parent (HP) and above high parent (AHP) than low parent (LP) and below low parent (BLP). SRCs with expression levels of AHP and HP tend to be 24-nt long, and SRCs with expression levels of BLP and LP tend to be 21-nt long. Between parents have 25% differentially expressed SRCs, and hybrid gain almost all in 22-nt SRCs and half in 24-nt SRCs. Expression level does not have global down or up tendency. However, public RNA-seq data show transposon is more inactivity in hybrid from 22-nt SRCs regions than their parents’. As for the miRNAs, 18 miRNA families were consisting downregulated in hybrid, but no one upregulated. *zma-miR528-5p*, *zma-miR408-5p*, *zma-miR1432-5p* were significant downregulated in hybrid, which may upregulate their target genes. Our results illustrated that sRNAs may contribute to heterosis at the very early stage of seed germination through repressing of retrotransposon activity, regulation gene activity in genic flanking regions, and promotion some gene expressions by downregulated miRNAs. Our data also can serve as a useful resource to better understand the sRNAs potential role in heterosis of maize.

## Results

### Characterization of sRNAs and SRCs in Maize

More than 120 million reads from eight samples were sequenced, and about 80% reads were perfect match on the maize B73 genome (Table S1). Maize have abnormal abundance of sRNA in 22-nt length, even more than the abundance of 21-nt sRNA, which mainly consist of miRNAs (Figure S1). In some varieties, the proportion of 22-nt sRNA is more than 24-nt sRNA’s (Figure S1). Maize sRNAs have more multiple loci on genome, nearly 50% of total reads have more than 20 loci (Figure S2A), and this ratio is less than 3% in *Arabidopsis thaliana* (Figure S2B).

Small RNAs prefer to consecutively express in genome, so we find high expressed small RNA clusters (SRCs), based on the mapping results. We found 84,689 SRCs across maize genome, which content 75.41% of total selected reads and coverage 0.56% of genome.

We can see the distribution of SRCs is similar with the distribution of genes (Figure 1A), and the SRC and gene number have positive correlation in each chromosome (Figure 1B). We classified the SRCs by their dominant length (e.g. If one cluster contain more than half 21-nt sRNA, then it classified as 21-nt SRCs), and found nearly 90% SRCs were 22/24-nt SRCs (Figure 1C). We distributed them on genome and found 22-nt SRCs especially enriched in repeat region (Figure 1D) and 24-nt SRCs enriched in repeat and genic flanking regions (Figure 1D/E). 20-nt and 21-nt SRCs usually have short length and high expression, miRNAs and ta-siRNAs are mainly in this group. Other SRCs contain a great number reads, because this group contain all the degraded products of tRNA and rRNA.

**Figure 1.**
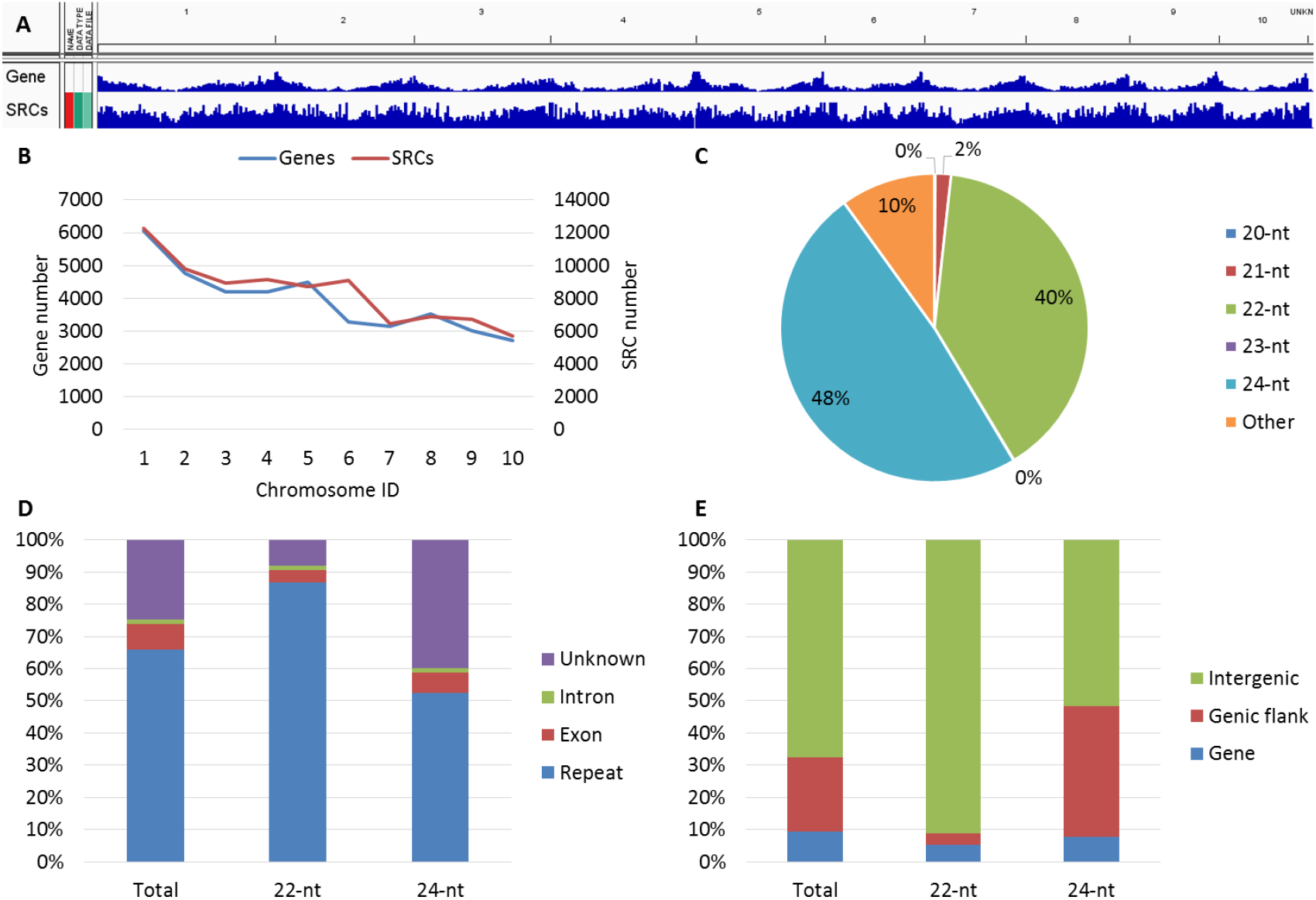
Summary and genome distribution of small RNA clusters (SRCs). A. SRCs and genes distribution on ten chromosomes; B. SRCs and genes number in each chromosome; C. SRCs classified by length of dominant (>50%) sRNA; D. Distribution of 22/24-nt SRCs on gene and repeat regions (Genes are classified into intron and exon; unknown means not annotated as gene or repeat regions); E. Distribution of 22/24-nt SRCs on gene and their flanking regions (±2 kb around genes).

### 22-nt SRCs enrichment in retrotransposon and 24-nt SRCs enrichment in DNA transposon

Here, we annotated these defined SRCs by using maize transposon element consortium (MTEC) annotation, and found the relationship between SRCs and repeat sequences. First, repeat sequences were separated into two types: retrotransposon, and DNA transposon. Then, we counted each type repeat and their region SRCs, and found the retrotransposon was account for more than 80% of total repeat, but only included less than 70% SRCs of repeat region, which mean DNA transposons were more activity than retrotransposons in originating SRCs (Figure 2A). We also checked the distribution of SRCs on retrotransposon and DNA transposon regions. Nearly 70% SRCs are 22-nt sRNA dominated in retrotransposon regions, and more than 80% SRCs are 24-nt sRNA dominated in DNA transposon regions (Figure 2B). 24-nt sRNA were widely studied^13, 15^, however, little known about potential functions of 22-nt sRNA in plant. We further focused on the 22-nt SRCs part, and found the 22-nt SRCs mainly from three super families of retrotransposon: Gypsy, Copia and Unknown, which all have less than 5% activity (Activity = SRCs numbers / repeat members * 100%) (Figure 2C). 22-nt SRC number and activity in each family were calculated, and we found gyma and ji families were the major contributor for 22-nt SRCs originating in Gypsy and Copia superfamily, but their activity vary from 3.1% to 14.6%. However, in some repeat families, such as raider, wiwa, and dagaf were had more than 30% activity in originating 22-nt SRCs (Figure 2D).

**Figure 2.**
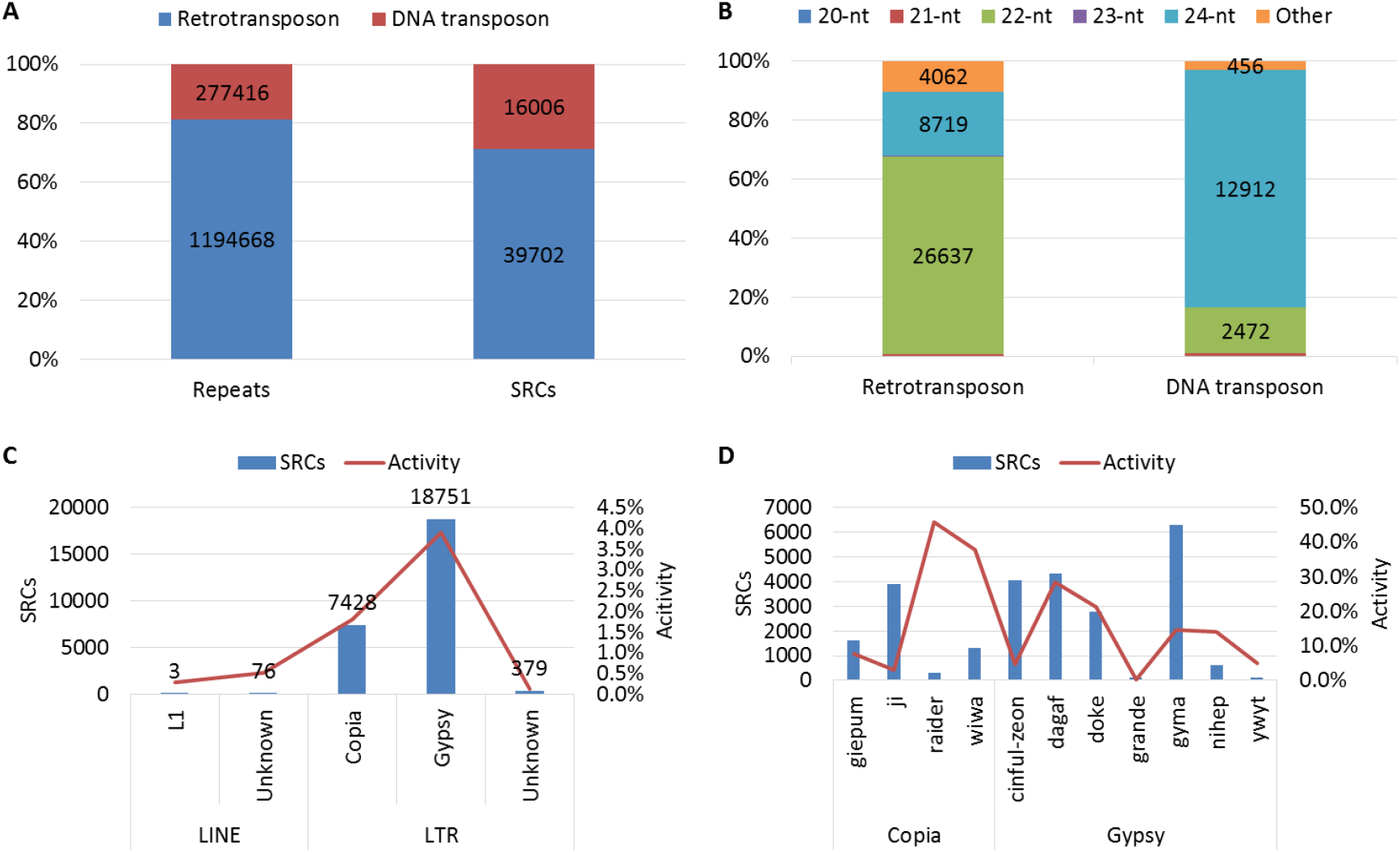
Distribution of SRCs on repeat regions. A. Distribution of SRCs on two types of repeat; B. Distribution of SRCs in different length on two types of repeat; C. Distribution of 22-nt SRCs on major super families of retrotransposon (Activity = SRCs numbers / repeat members); D. Distribution of 22-nt SRCs on major families of retrotransposon

### Expression of SRCs among parents and hybrid

To show expression patterns of SRCs in hybrid, we classify the SRCs into 7 groups according the expression among hybrid and parents (detail in method part). We showed genomic distribution and expression pattern of SRCs, and found most of SRCs were middle parent (MP) expression in hybrid (Figure 3). Another apparently expression pattern of SRCs in high parent (HP) and above high parent (AHP) were more than low parent (LP) and below low parent (BLP) (Figure 3). This expression pattern conserved in three different cross combinations, especially in 24-nt SRCs region (Figure 3, Figure S3A/B). However, the 22-nt SRCs have different patterns in each cross combination (Figure 2, Figure S3A/B). We further compare three cross combinations, and found some SRCs had conserved expression patterns in different hybrids. An interesting result showed nearly all common HP and/or AHP SRCs were 24-nt SRCs, and common LP and/or BLP SRCs were 21-nt SRCs (Figure 4).

**Figure 3.**
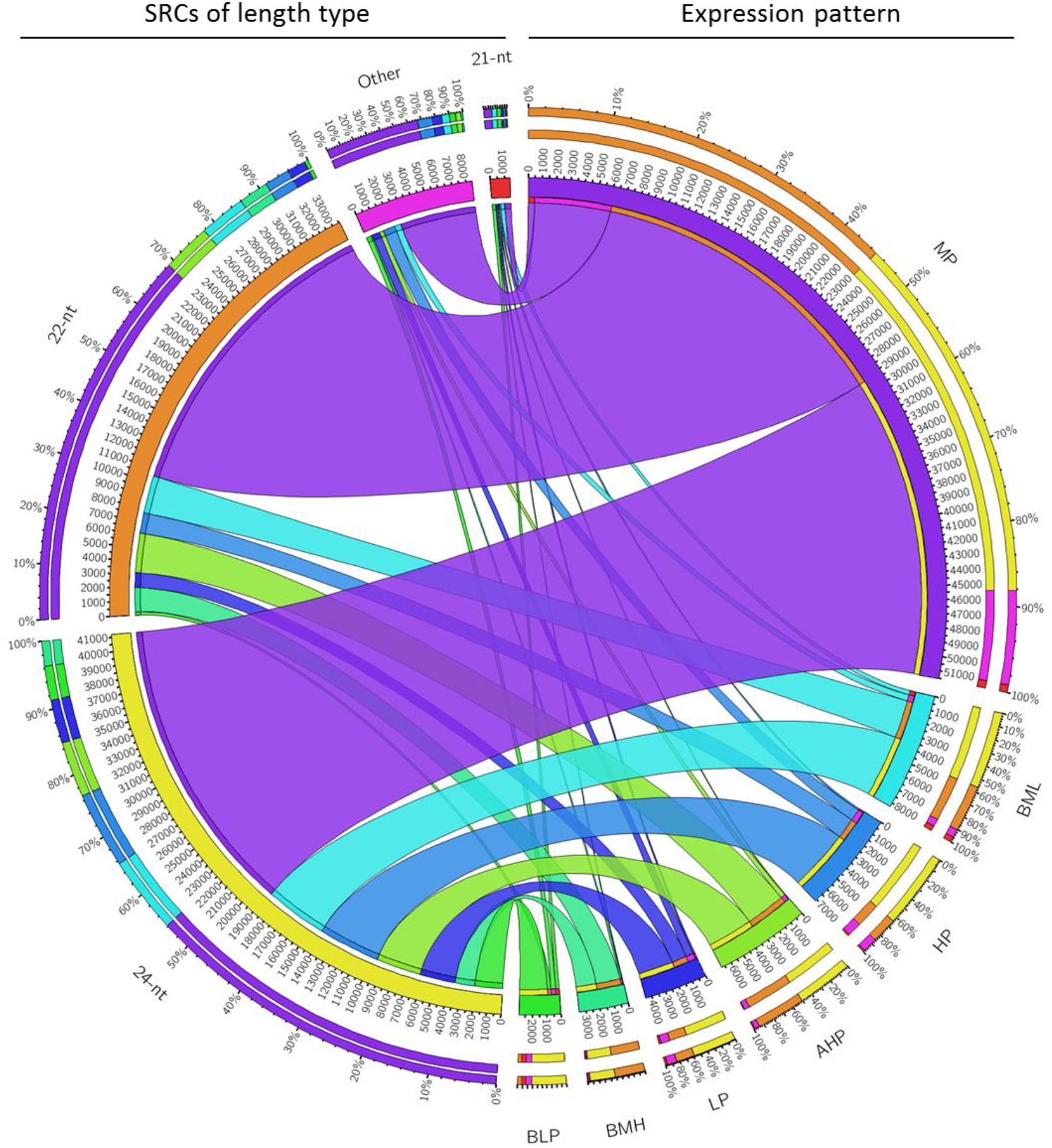
Classified SRCs based on expression between parents and hybrid. (SRCs are group into 7 classes, from high to low are above high parent (AHP), high parent (HP), between MP and HP (BMH), middle parent (MP), between MP and LP (BML), low parent (LP), and below low parent (BLP).)

**Figure 4.**
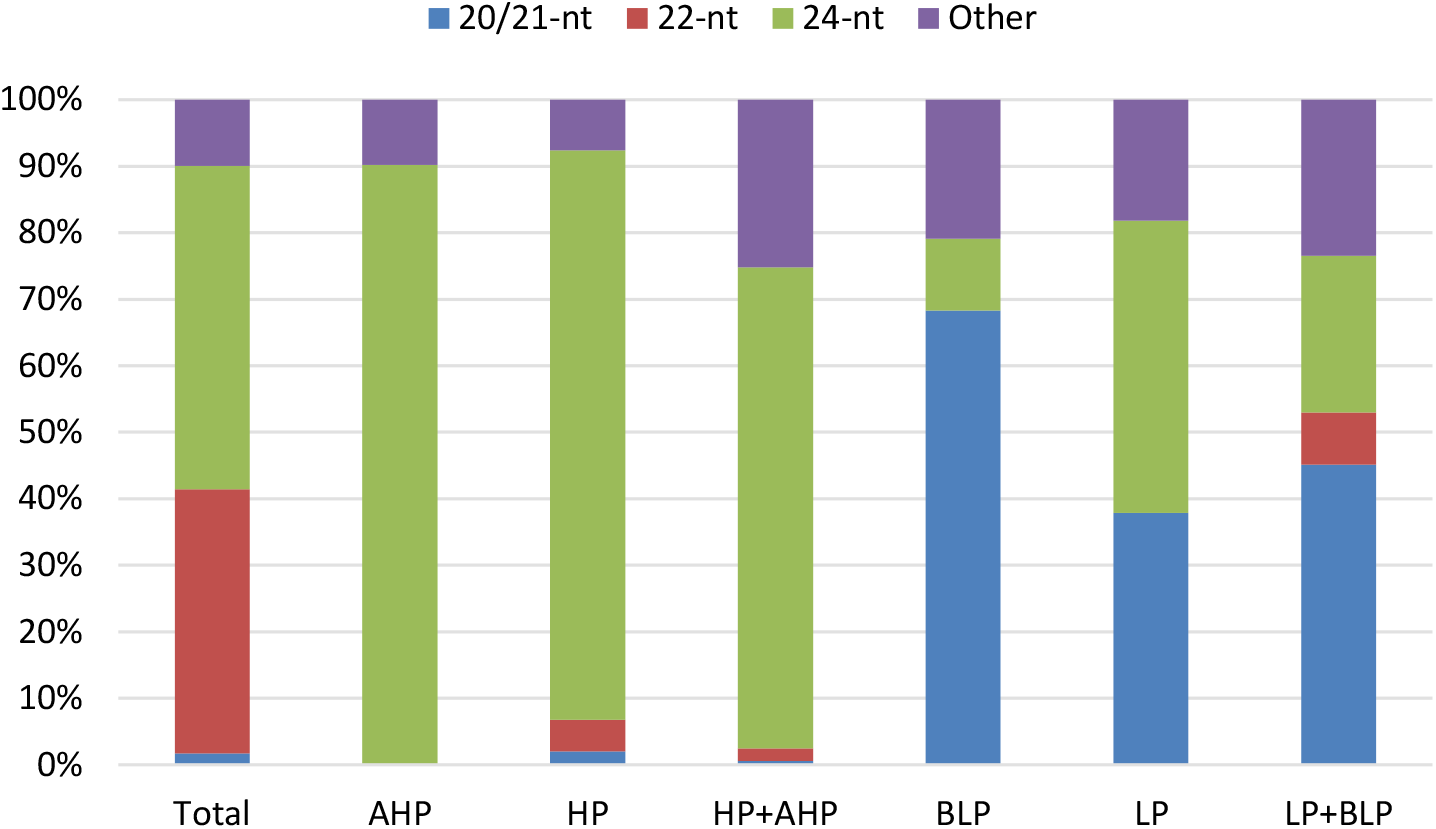
Proportion of each length type SRCs in conserved expression pattern. AHP & HP almost all 24-nt SRCs, and BLP and LP enrichment in 21-nt SRCs.

Above mentioned 24-nt SRCs enriched in genic flanking regions (Figure 1E), and some of them had conserved HP and/or AHP expression patterns (Figure 4). So, we further discussed these 24-nt SRCs relationship with genes, and found 24-nt SRCs great enriched in 1 kb region of gene flank (Figure 5A). These genes, which include 24-nt SRCs in their flank (1 kb) regions, were enrichment in cellular process and metabolic process (Figure 5B). And these SRCs expression of hybrids was higher than their parents’ in each biological process (Figure 5C/D).

**Figure 5.**
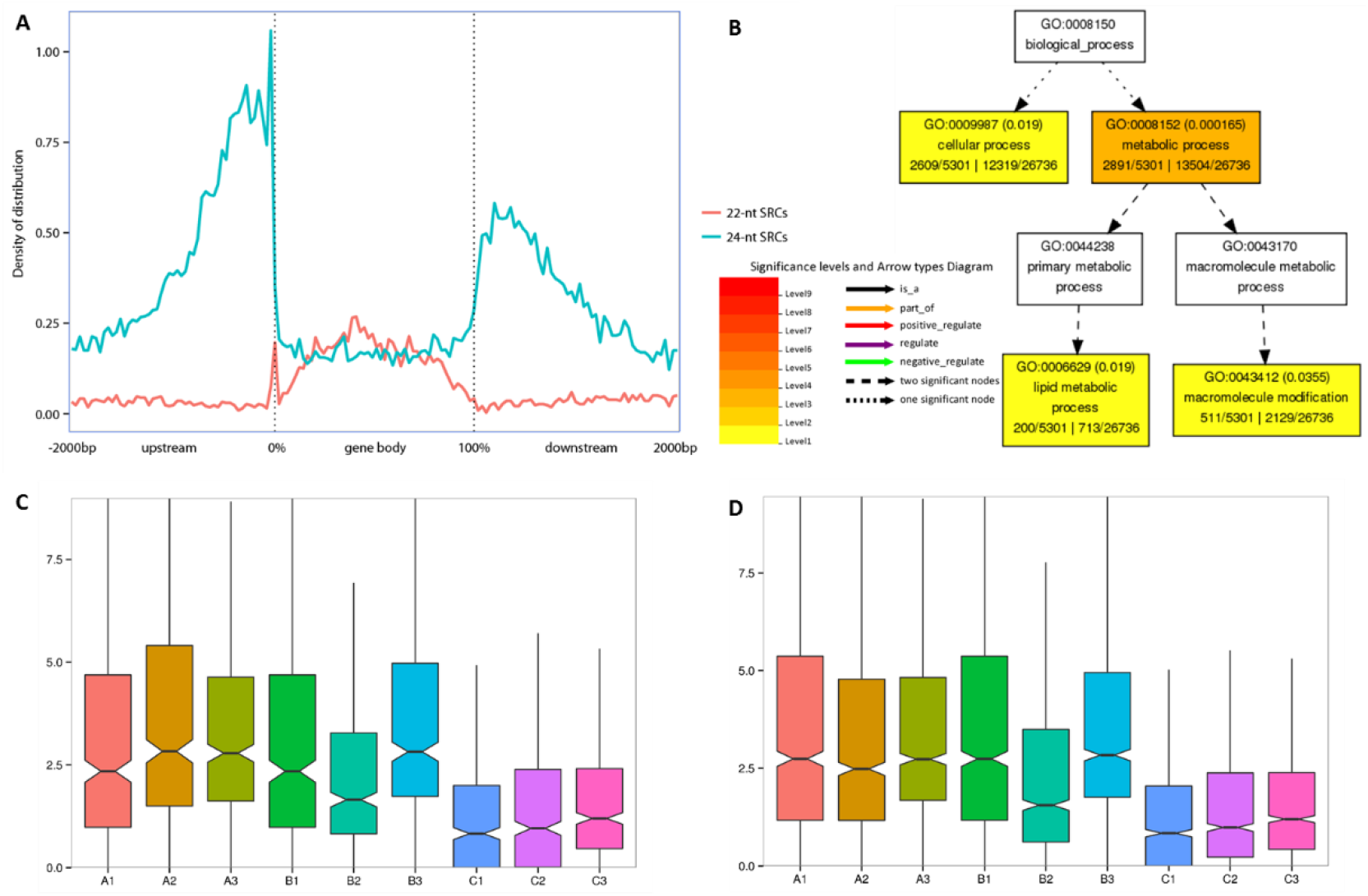
24-nt SRCs enriched in gene flank and preference upregulated in hybrid. A. Distribution of 22/24-nt SRCs on gene bodys and their flanking regions (All the gene body scale to 0 to 100%, and flanking regions include 2 kb of upstream and downstream of gene. Each part (upstream, gene body, and downstream) split into 50 bins, and density is SRCs number in each bin). B. Enrichment GO terms of genes, which include 24-nt SRCs in their flank 1 kb region. C. 24-nt SRCs expression in flank regions of lipid metabolic process related gene, A/B/C represent each cross combination. D. 24-nt SRCs expression in flank regions of macromolecule modification related gene.

### Parents exist great difference in SRC regions and inheritance by hybrid

The percentage of 2-fold different SRCs between parents was very high, nearly a quarter of them have more than twice difference in expression (Figure 6A). We found 3258 SRCs had twice difference, and common exist in 3 cross combinations (Figure 6B), but no enrichment or bias distribution of these SRCs were found.

**Figure 6.**
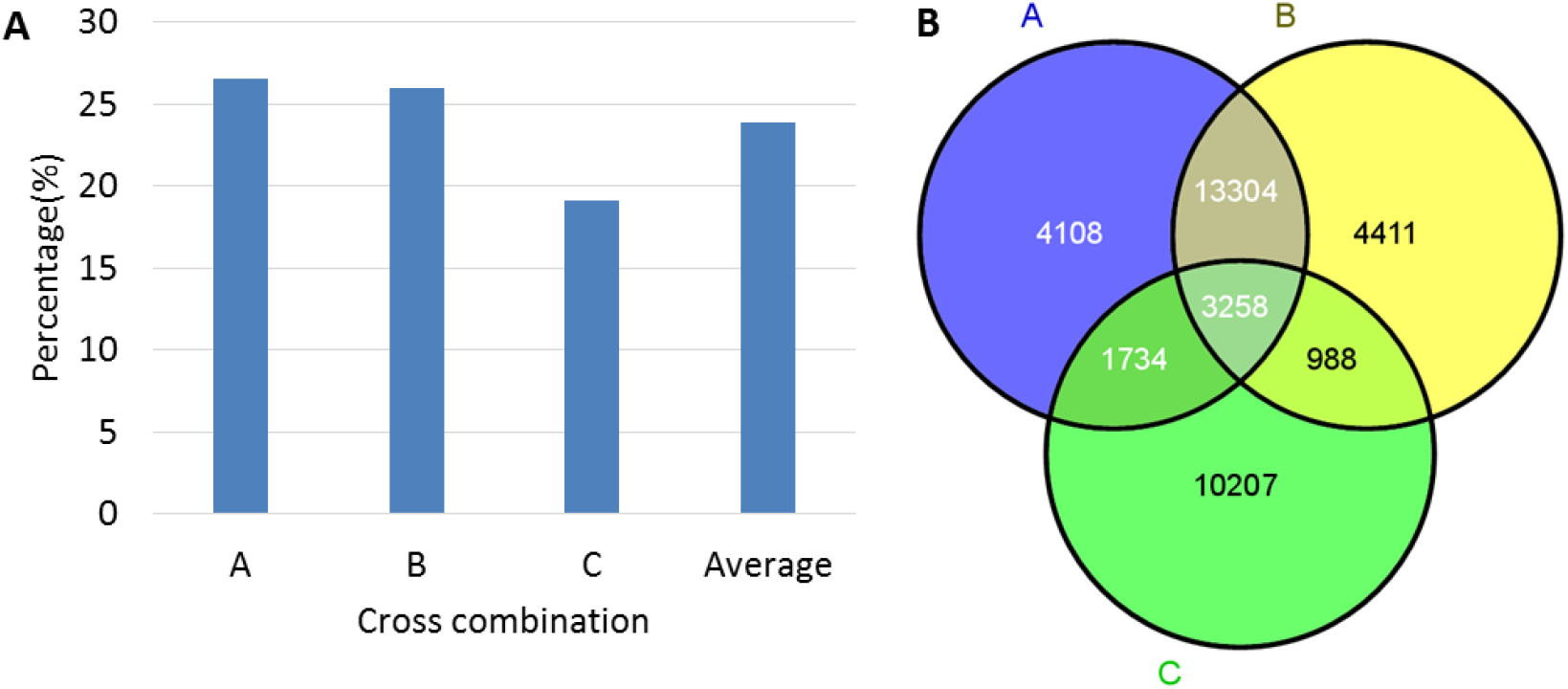
2-fold different SRCs between parents (A) percentage in each cross combination, (B) Overlapping among 3 cross combinations.

To investigate the different SRCs between parents and hybrid, reads per million (RPM) more than 5 was used to filter highly expressed SRCs in each sample. We found many variant highly expressed SRCs between parents in 22/24-nt SRCs (Figure 7 A/B). About 21% of 22-nt highly expressed SRCs were different between paternal and maternal, and hybrid gain almost all different SRCs (Figure 7A). This pattern was also conserved in other two cross combinations (Figure S4 A/E). However, the expression tendency between hybrid and parents were not found in these 22-nt SRCs (Figure 6C). As for the 24-nt highly expressed SRCs, only a half of parents specific SRCs were gained by their hybrid (Figure 4B). The similar pattern also shows in B and C combinations (Figure S4 B/F). The expression tendency between hybrid and parents were not found in these 24-nt SRCs of A cross combination (Figure 7C/D), but have a slight downregulated in B and C cross combination (Figure S4 C/D/G/H).

**Figure 7.**
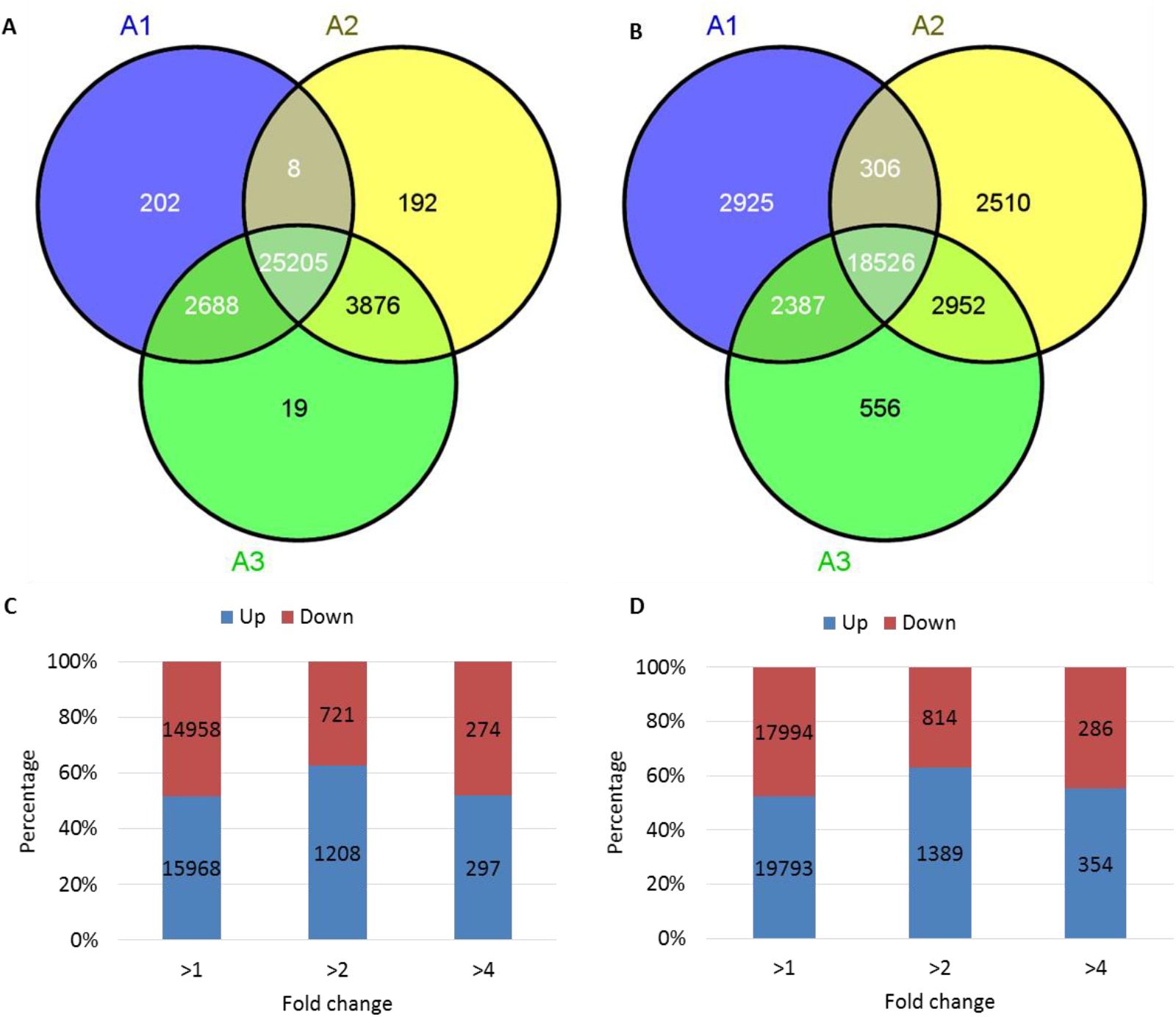
Expressed SRCs among parents and hybrid in A cross combination. (A). 22-nt SRCs of parent specific were good inherited in hybrid. (B). 24-nt SRCs of parent specific were half inherited in hybrid. (C). Differentially expression 22-nt SRCs under different threshold. (D). Differentially expression 24-nt SRCs under different threshold.

### miRNAs are downregulated in hybrids

Compare with middle parents’ values (MPVs), percentage of miRNA show a tendency of downregulated in hybrids. This pattern was conserved in three cross combinations (Figure 8). Percentage of miRNA even lower than their both parents in B and C combinations.

**Figure 8.**
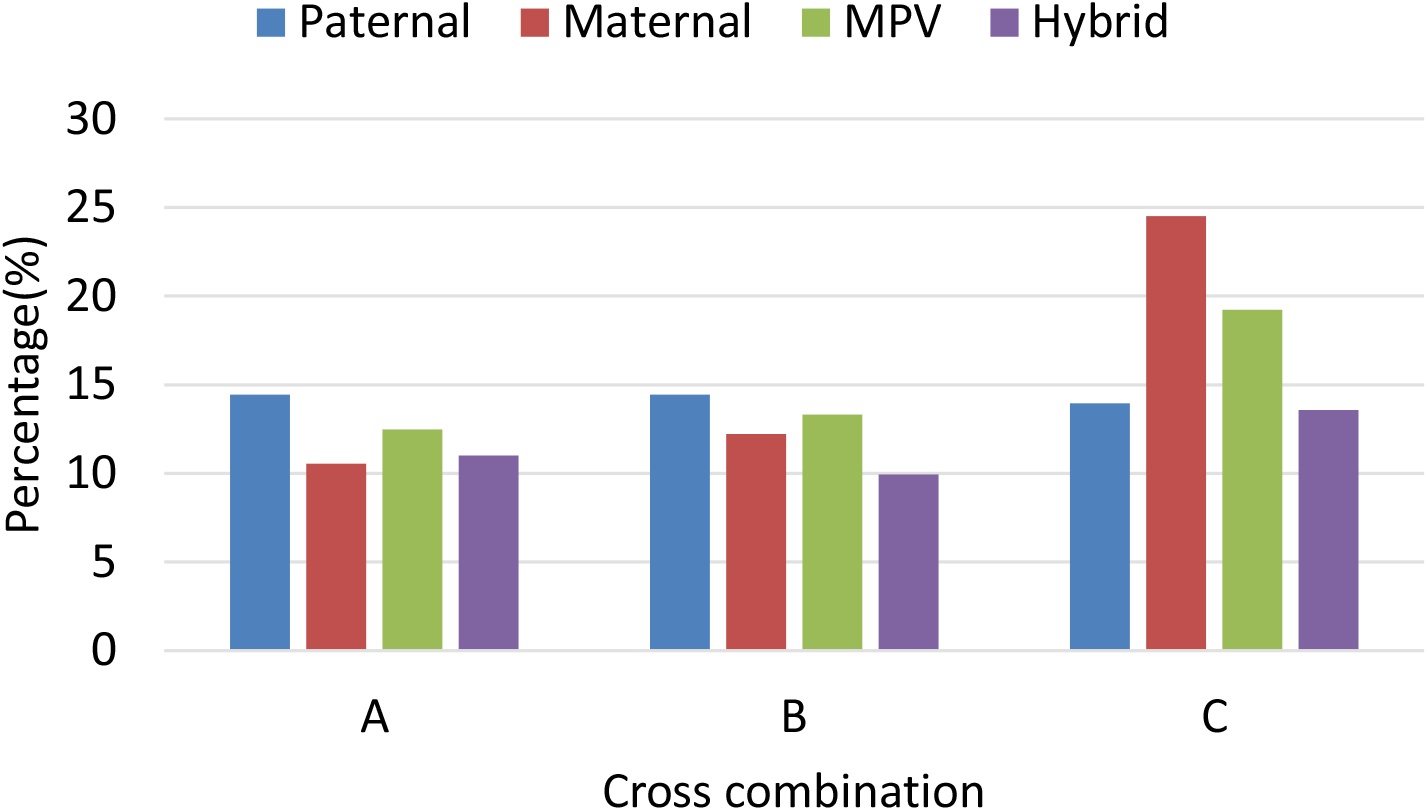
Percentage of miRNA in each sample and tendency downregulated in hybrid

To investigate which miRNAs changed in hybrids, we counted the RPM values of each miRNA family and showed their expression patterns (Figure 9). The results showed that most of miRNA families were downregulated in hybrid, compared with MPV (Figure 9). B and C combinations were also show the similar pattern (Figure S5). Three cross combination results show a conserved pattern that miRNA global downregulated in hybrids.

**Figure 9.**
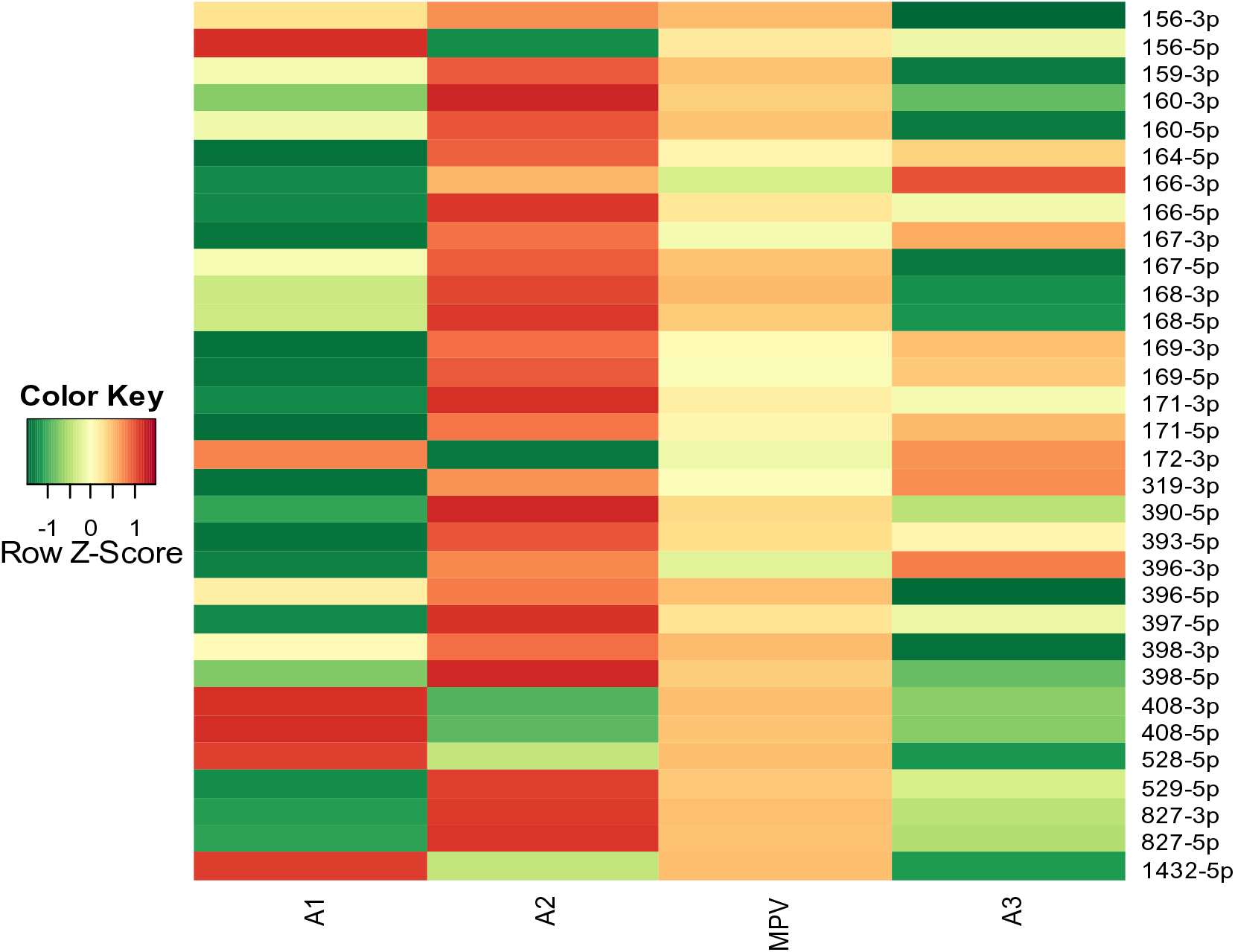
Known miRNA family mature expression in A cross combination, all the miRNA ID omit zma-miR (normalize to Z-Score)

As for 32 miRNA families, which expression more than 5 RPM, we found 18 of them were common downregulated in three combinations. Among of them, 2 miRNA families (*zma-miR408-5p, zma-miR1432-5p*) are 2-fold downregulated, and 6 miRNA families (*zma-miR528-5p, zma-miR408-3p, zma-miR168-3p, zma-miR156-3p, zma-miR529-5p, zma-miR398-5p*) are 1.5-fold downregulated. As well as, none miRNA family was common upregulated in three combinations.

## Discussion

Maize have a big part of 22-nt sRNA originate from retrotransposon, and even higher expressed than 21-nt or 24-nt sRNA in some varieties (Figure S1A/B/C). We also compared with Arabidopsis thaliana sRNA data, and found maize sRNA of 21 and 24-nt length decreased was due to dilution of increased 22-nt sRNAs (Data download from GEO, Figure S1D).

### 22-nt SRCs gained by hybrid may better repress retrotransposon expression

22-nt sRNA is great enriched in maize and lack in Arabidopsis (Figure S1), but 22-nt SRC is the second large (40%) group in maize (Figure 1C). However, the function of 22-nt SRCs is seldom known in maize. Many papers reported 24-nt siRNAs suppress DNA transposon expression through RNA directed DNA methylation (RdDM) pathway ^12, 13, 16, 17^. In this study, we found nearly 70% 22-nt SRCs enriched in retrotransposon regions (Figure 2B). Does these 22-nt SRCs have relationship with retrotransposon, and have conserved pattern in hybrids? Above result showed 22-nt SRCs do not show a consistency tendency of upregulated or downregulated in hybrid (Figure 7C, Figure S4C/G), but the kinds of expressed 22-nt SRCs were great abundance in hybrid, which inherited nearly all expressed 22-nt SRCs in parents (Figure 7A, Figure S4A/E). To prove whether variety 22-nt SRCs could influence retrotransposon in hybrid, we downloaded public RNA sequencing (RNA-seq) data about maize hybrid and their parents^18^, and analysis the expression pattern of DNA transposon and retrotransposon on SRC regions. We found the expression of retrotransposon, which originated of 22-nt SRCs, was downregulated in hybrid, and this pattern conserved in root and shoot (Figure 10A/B). The tendency of downregulated transposons was only found in 22-nt SRCs regions, 24-nt SRCs regions were not have the same pattern (Figure S6). This result gives a clue, hybrid gain diversity 22-nt SRCs may better control activity of retrotransposons, reduce waste energy and materials on transposon, maintain integrity of genome. All of these potential results will benefit to hybrids and show strong phenotype in hybrid vigor.

**Figure 10.**
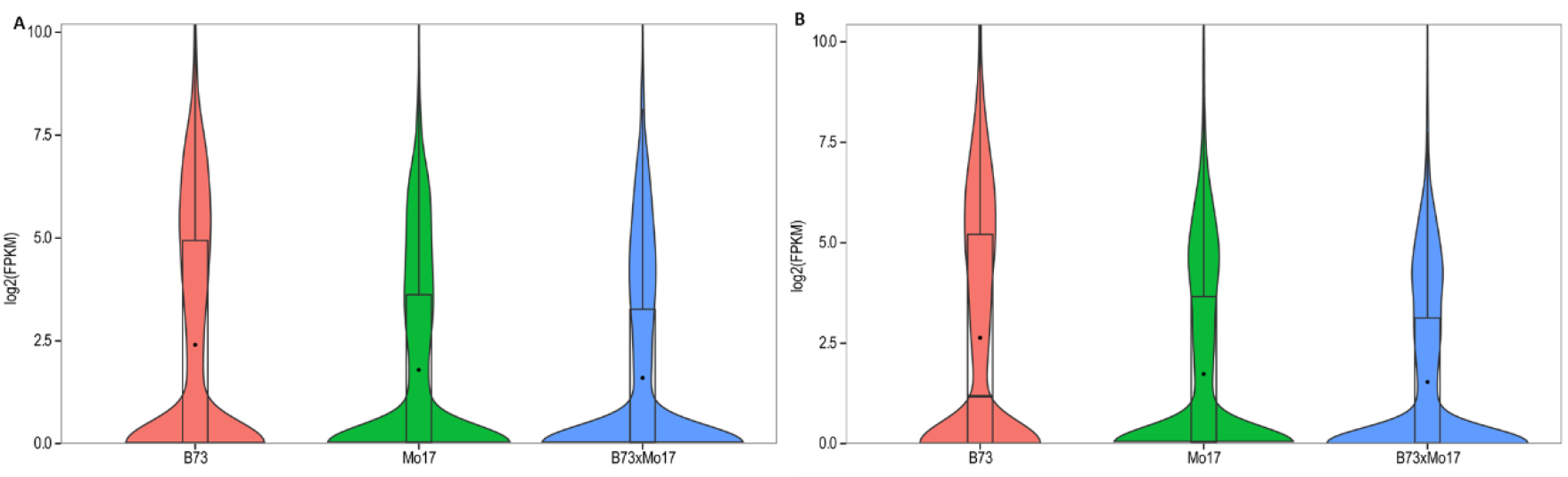
Expression of retrotransposon downregulated in hybrid from 22-nt SRC regions. (A) root, (B) shoot.

miRNAs downregulate may cause some genes increased and contributed to hybrid vigor. Ding, et al.^6^ reported global downregulated of miRNA in hybrid. In this study, we also found the similar pattern in three hybrids. To elusive the potential functions of conserved and downregulated miRNAs, the miRNA targets were predicted by psRobot online^19^. *Zma-miR408-3p/528-5p* are predicted co-targeted on LAC2 genes (*GRMZM2G367668 and GRMZM2G169033*), which have conserved target site in some dicots (include *Sorghum bicolor*, *Oryza sativa*, *Brachypodium distachyon*), and show shorten elongation of root in knockout mutation ^20^. *Zma-miR408-3p* also predicted targeted on SVR7 gene, which is a kind of pentatricopeptide (PPR) repeat-containing protein accumulation and translation of chloroplast ATP synthase subunits. Relief repress of these genes may promote elongation of root and accumulation more chloroplast ATP synthase for photosynthesis, in order to show strong hybrid vigor in hybrids.

## Method

### Plant materials, RNA extraction and deep sequencing

To investigate whether small RNAs involved in heterosis, and a conserved expression pattern of sRNAs exist in different cross combinations, next generation deep sequencing technique was performed. In this study, the three hybrid variants of maize, which were elite hybrids and widely cultivated in China that exhibited high heterosis for grain yield, and theirs five parental inbred lines were utilized (all samples relations were show in the figure 11). Fifteen seeds of each hybrid and parental line were surface-sterilized in 70% (v/v) ethanol for 10 min, and then rinsed several times with sterile distilled water. Then the seed were soaked in sterile distilled water and incubated at 26°C in the dark for 24 hours. The embryo tissues were peeled from the seeds, and each embryo tissue material from same variety was mixed for RNA extraction. Total RNA from the all samples was extracted with TRIzol reagent (Invitrogen, 15596-026) following the manufacturer’s protocol. Each sample of 15 ng RNA was used for small RNA sequencing libraries prepared and on Illumina HiSeq2000 system by BGI.

**Figure 11.**
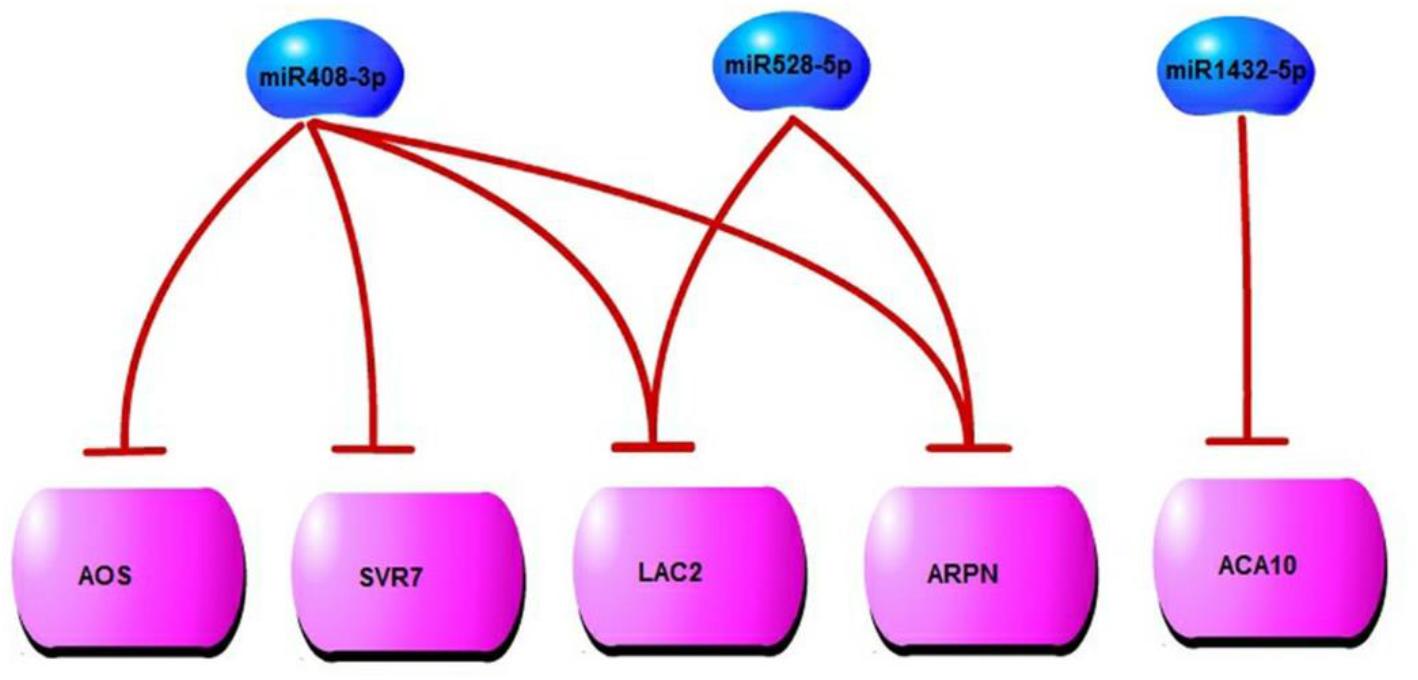
miRNA downregulated may contribute to heterosis in maize embryo

**Figure 12.**
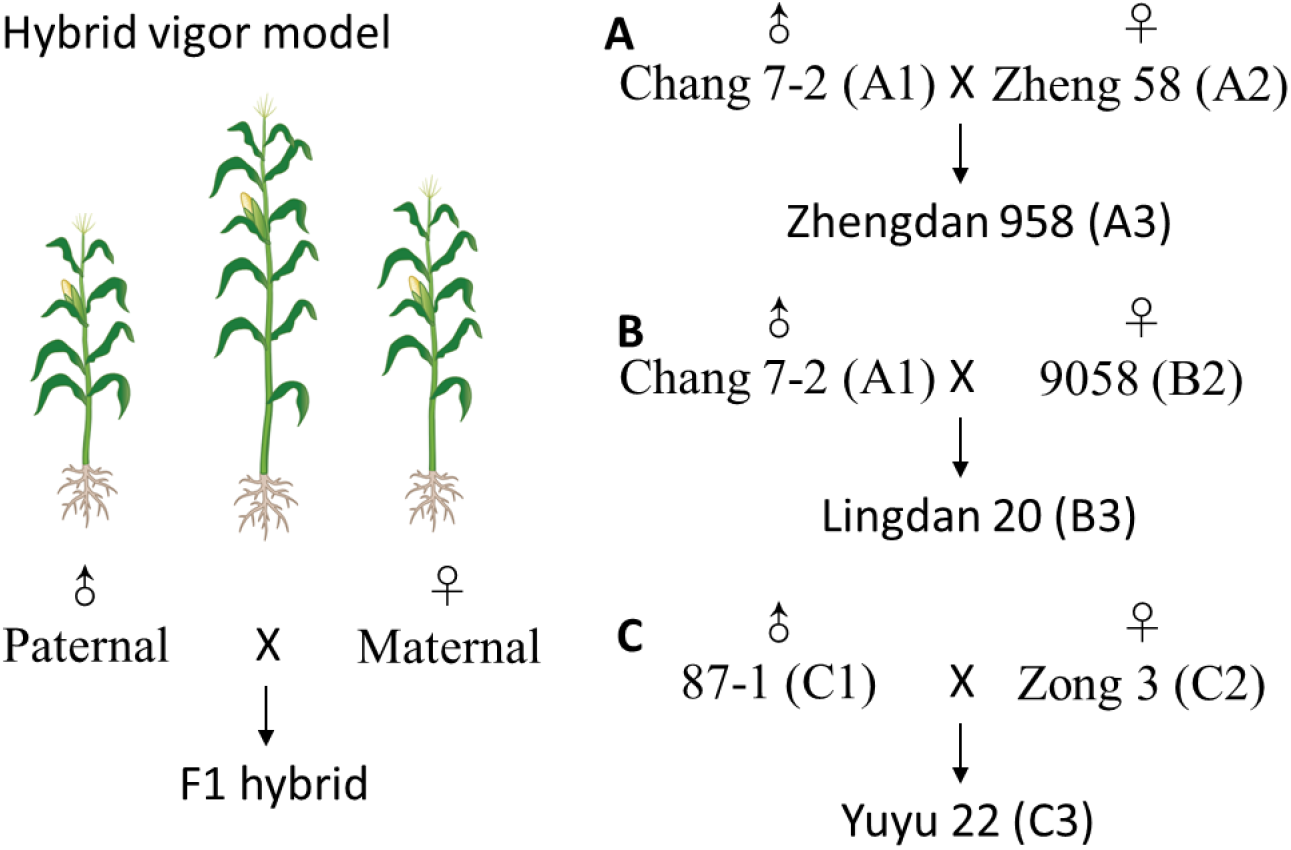
Used varieties of maize, A, B, C represent three cross combinations.

**Figure 13.**
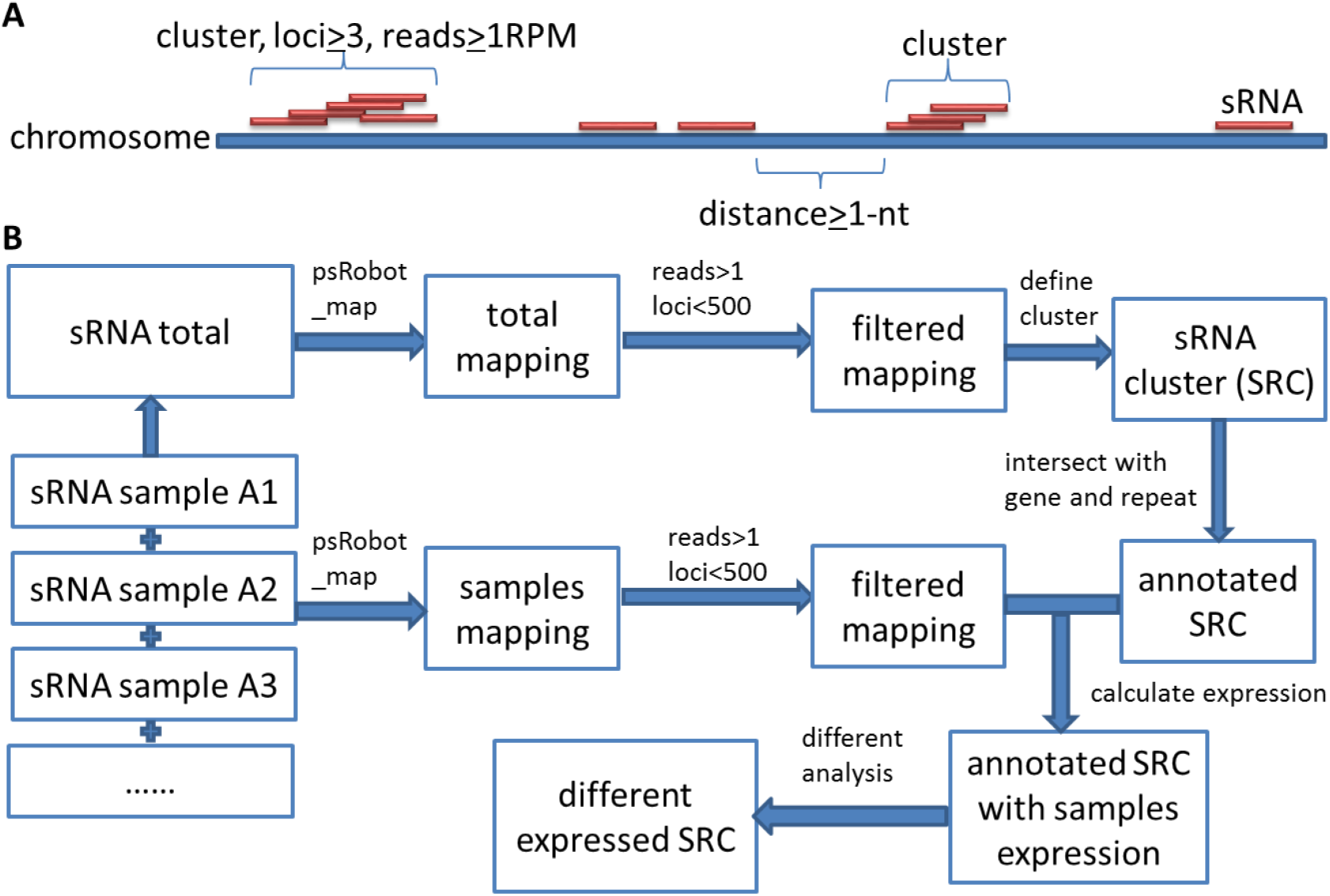
Define and analysis small RNA clusters (SRCs). (A) Model of define SRCs; (B) Pipeline for analysis SRCs.

**Figure 14.**
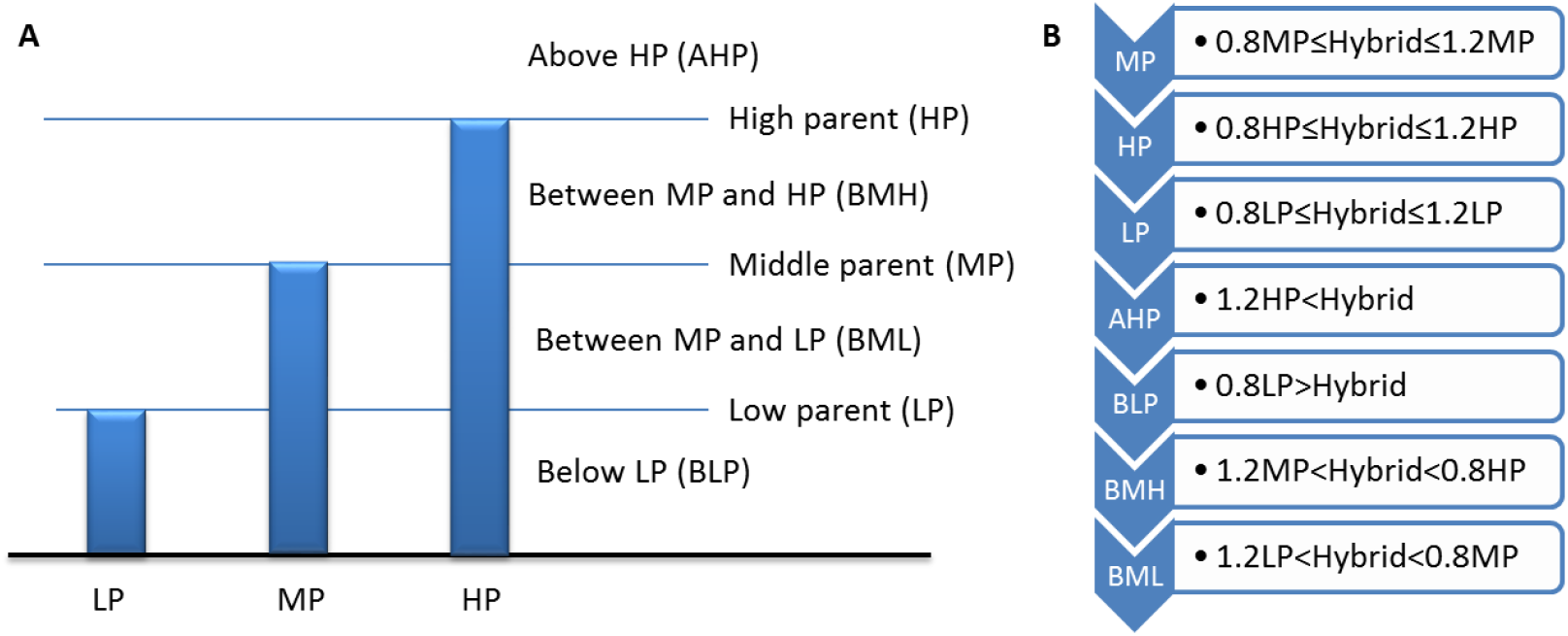
Flow chart for classifying SRCs based on hybrid expression compared to parents’ Based on above procedure, we get each SRC expression pattern, and show theirs relation with SRCs’ length type distribution by Circos^25^.

### Identification small RNA clusters and analysis difference

Maize genome were download from Maizesequence, and filtered gene set (FGS) and TE Consortium repeat annotations (MTEC) were used^3^. Small RNAs (sRNAs) were mapping on the reference genome by psRobot toolkits^19^. Maize gene annotations of homolog were downloaded from phytozome^21^. Maize tRNA and rRNA were identified by RepeatMasker 4.01 (www.repeatmasker.org), based on RepBase version 20130422 ^22^. Based on all the sRNA mapping result, highly and consecutive expressed sRNA region were identified as sRNA clusters expressed the following criteria (following figure): (1) Sequence need to perfect match reference genome; (2) Select loci (hits) < 500 mapping sequences for exclude highly repeat sequences; (3) Select reads counts >1 sequences in order to exclude occasion transcription or sequencing noise; (4) sRNA consecutive detected in one cluster, any cluster distance >1 nt as independence cluster. (5) In each cluster contain loci number ≥ 3, read per million (RPM) value ≥1, and reads per million (RPM) value ≥ 5.

Statistics of SRCs basic information were procedure by Perl script, and histogram were draw by Microsoft Excel 2013. sRNA mapping result compare with SRCs were done by intersectBed from BEDtools 2.16.2^23^. Distribution of SRCs and genes were showed by IGV 2.2.4^24^.

SRCs have highly correlation with repeat, and two types of repeat annotations exist in maize genome: maize transposon element consortium (MTEC) and MIPS/Recat. We selected MTEC for further study, due to it have more TE members, detail annotation and classification than MIPS. As for the 1971471 annotations of MTEC, we found some information is redundancy, such as same loci have duplication annotations, similar repeat members of same family overlap each other. After remove redundancy and merge overlapped repeat in same family, we got 1472084 repeat annotations. Compare annotations with SRCs by BEDtools ^23^, and result summary by Perl script.

To explore expression pattern of 22-nt SRCs in retrotransposon (RT22-SRCs), we compare them between hybrid and middle parent (MP), and select fold change > 2, Fisher test P-value <0.01 and false discovery rate (FDR) <0.05 by Perl scripts.

For discussion expression pattern of SRCs between hybrid and parents, we classify the SRCs into 7 groups based on SRCs expression of hybrid relation with parents’, include high parent (HP), low parent (LP), average expression between parents called middle parent (MP), above high parent (AHP), below low parent (BLP), between HP and MP (BHM), and between HP and LP (BHL). The classify procedure is in the following figure.

### Transposon expression on SRC regions

To show whether SRCs regulated transposon, we downloaded RNA-seq data of maize hybrids and their parents from NCBI(GEO ID: GSE43142)^26^, include 6 samples of inbred lines Mo17 and B73, and their hybrid in shoot and root. Firstly, transformed SRA data to fastq format by fastq-dump from SRA Toolkit (Version: 2.1.12, http://eutils.ncbi.nih.gov/Traces/sra/?view=software). Secondly, removed low quality bases and shot reads by fastq_quality_trimmer from fastx toolkit (Version: 0.0.13, http://hannonlab.cshl.edu/fastx_toolkit/), parameters in -t 20 (trim phred score of base lower than 20 from the reads end) and -l 20 (remove length of trimmed reads lower than 20-nt), and then check the quality of data by FastQC (Version: 0.10.1, http://www.bioinformatics.babraham.ac.uk/projects/fastqc/). Thirdly, created index of maize genome by bowtie-build, and then using bowtie (Version: 0.12.7, parameter used -v 2 -m 500 -S-p 5) mapped reads of RNA-seq to reference genome^27^. Then, filtered unmapped reads and transformed to BAM format by SAMtools (Version: 0.1.18)^28^. Finally, the FPKM values of transposon in SRC regions were calculated by cuffdiff from cufflink (Version: 2.1.1)^29^. The FPKM values in each sample are showed in violin plot by R language.

### miRNA expression and their target genes

All the sRNAs were mapped on known pre-miRNA of maize from miRbase 19.0 by psRobot local version 1.01^19, 30^. The percentage of miRNA in total sRNA was calculated by Perl scripts. All the miRNA expressions were normalized to reads per million (RPM), and grouped by family. The expression of each miRNA family was showed in heatmap and normalized to Z-score by R language. For the potential targets of miRNA, and conservation of target sites were all predicted by psRobot online^19^.

## Acknowledge

The authors thank Dr. Jihua Tang for kindly providing maize inbred line 87-1. The authors also thank Dr. Hua-Jun Wu and Meng Wang for the suggestion of data analysis.

## Supplementary

**Table S1.** Statistics of sRNA sequencing and mapping

| ID | Variety | Total reads | Mapping reads %<sup>1</sup> | Selected reads %<sup>2</sup> |
| --- | --- | --- | --- | --- |
| A1 | Change 7-2 | 16966913 | 74.32% | 68.78% |
| A2 | Zheng 58 | 15810440 | 77.77% | 62.57% |
| A3 | Zhengdan 958 | 18399480 | 76.55% | 64.27% |
| B2 | 9058 | 18590068 | 80.71% | 67.91% |
| B3 | Lingdan 20 | 17249792 | 76.38% | 64.96% |
| C1 | 87-1 | 9735651 | 83.62% | 51.77% |
| C2 | Zong 3 | 13150801 | 83.55% | 50.58% |
| C3 | Yuyu 22 | 11068163 | 84.00% | 44.63% |
| Merge |  | 120971308 | 79.02% | 66.90% |

<sup>1</sup> Mapping reads%=Perfect matched reads/Total reads; <sup>2</sup> Selected reads%= Selected reads/ Perfect matched reads, selected reads need reads count > 1, reads loci <500 and not mapping on tRNA and rRNA region.

**Figure S1.**
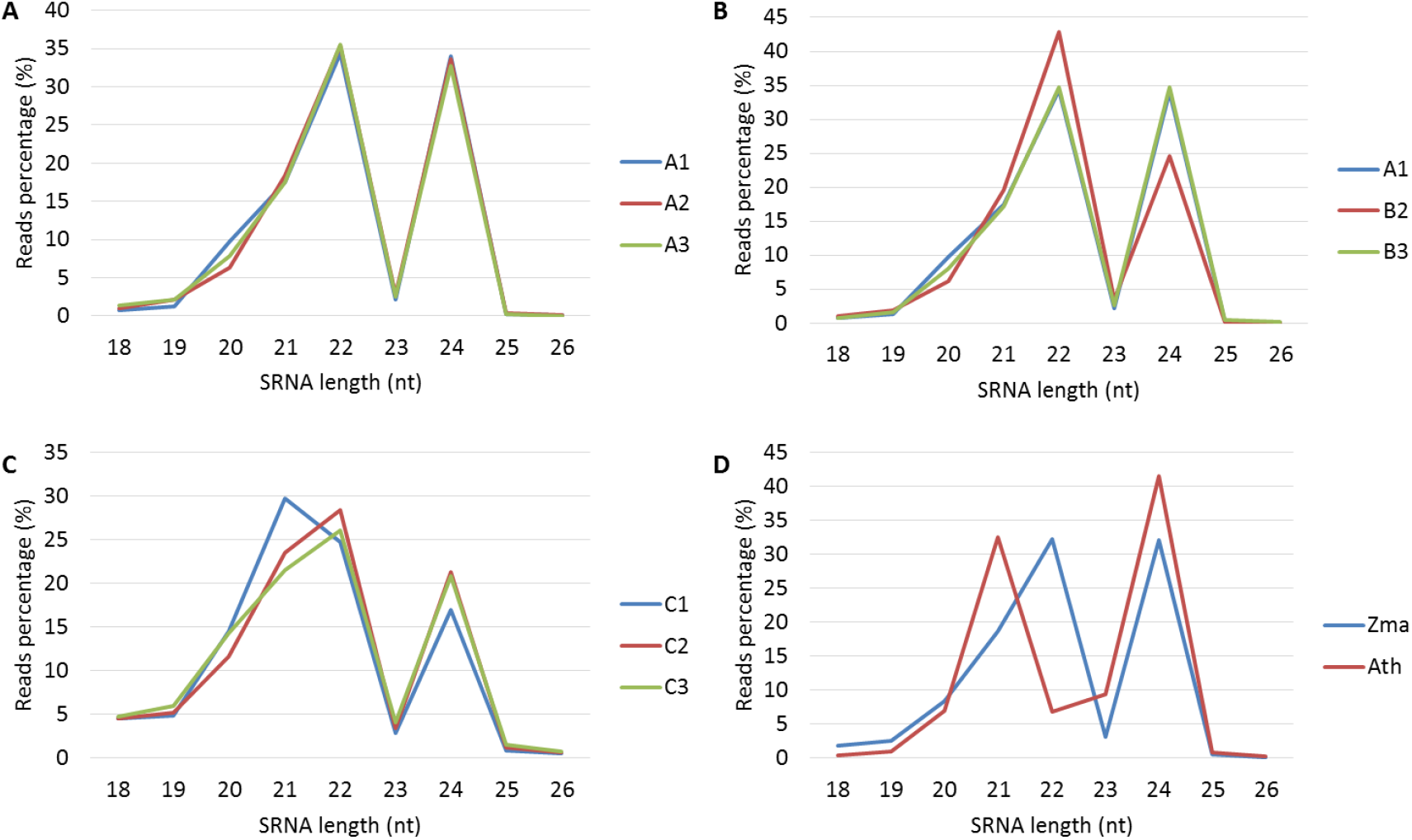
Summary of sRNA total reads length distribution (A/B/C) sRNA length distribution in A/B/C cross combination, (D) sRNA length distribution of *Zea may* and *Arabidopsis thaliana*.

**Figure S2.**
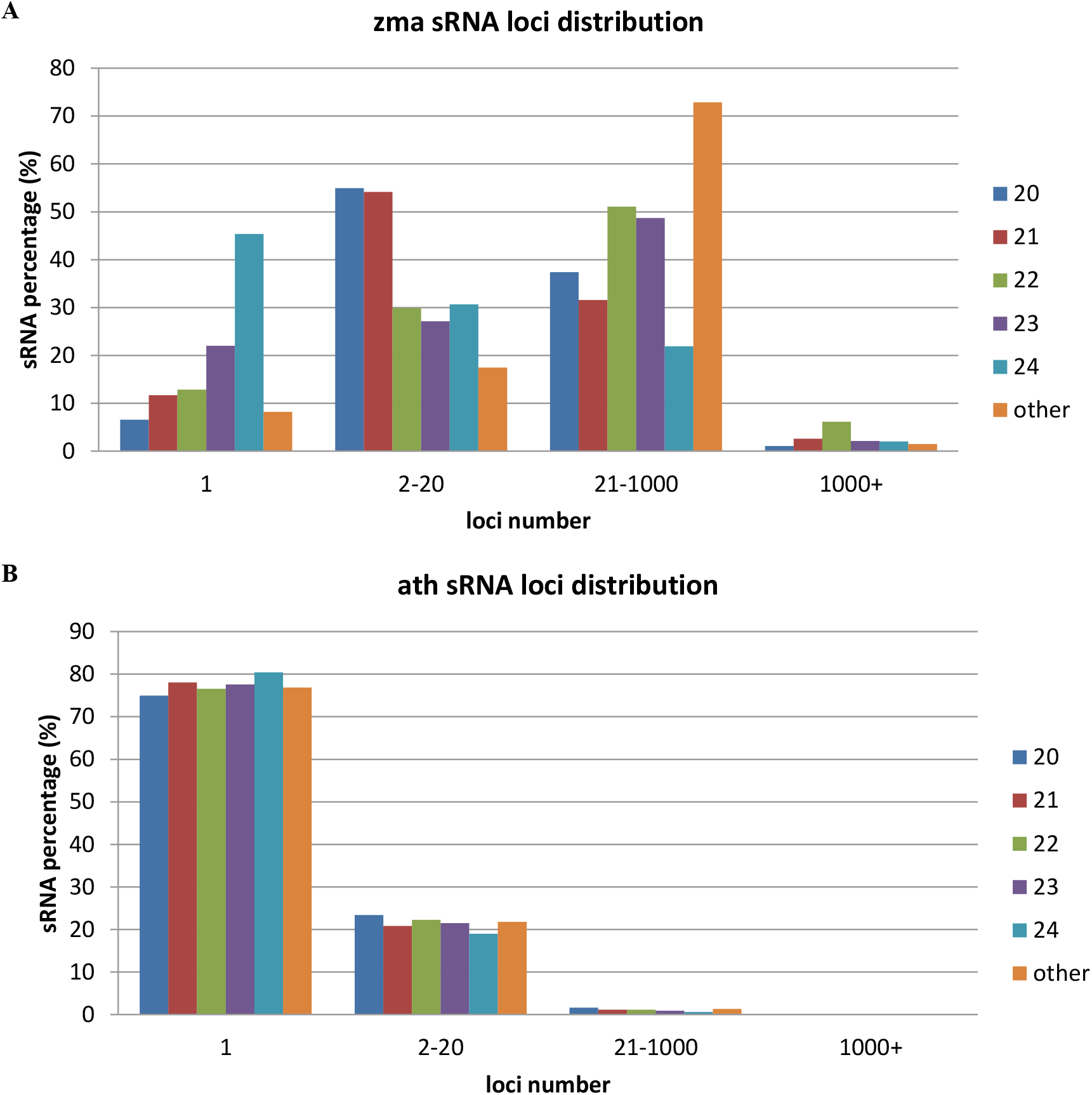
Different length total sRNA loci and distribution, (A) Zea mays, and (B) Arabidopsis thaliana. 94.87% sRNAs in Arabidopsis thaliana less than 20 loci. However, sRNA in maize have very high percent repeat, more than half of them reads have more than 20 loci. If use loci threshold 20 will exclude nearly 43.4% mapped reads. But use threshold of 1000, only 3.2% mapped reads were excluded.

**Figure S3.**
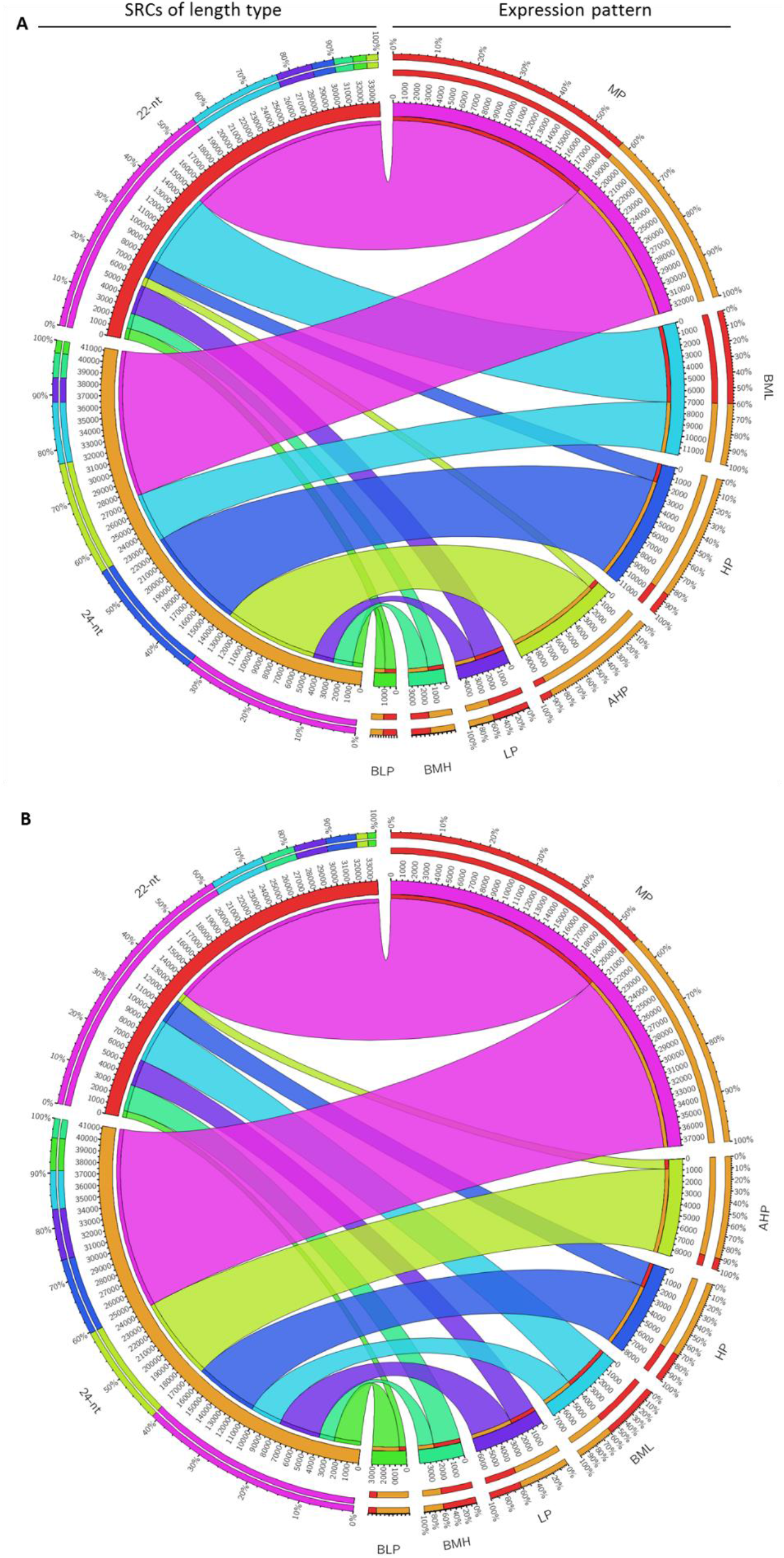
Compared with parents, length type SRCs expression changed in hybrid. (A) B cross combination; (B) C cross combination.

**Figure S4.**
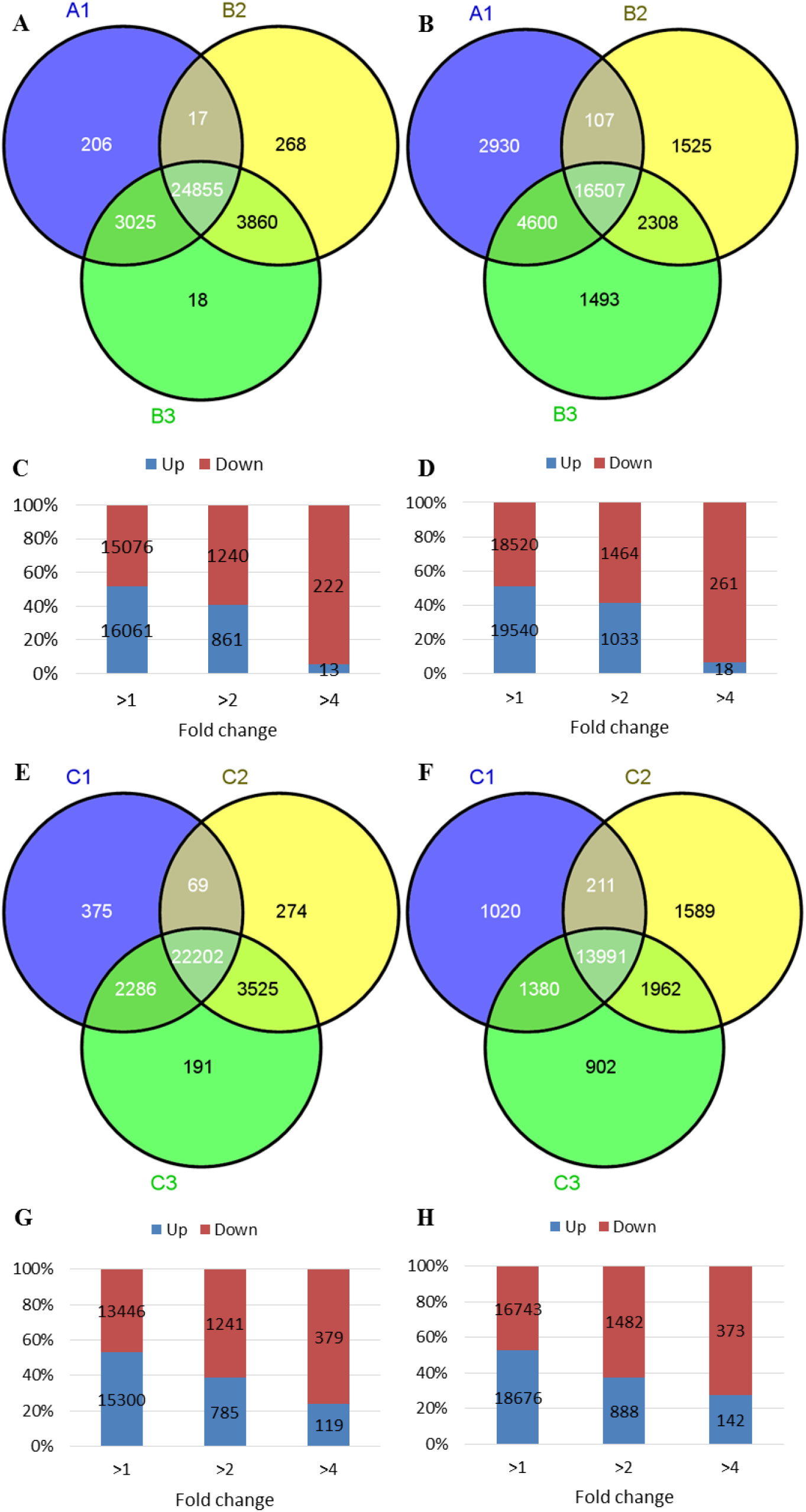
SRCs between parents and hybrid in B and C cross combination. (A-D) from B combination, (A/C) 22-nt SRCs expression; (B/D) 24-nt SRCs expression. (E-H) from C combination, (E/G) 22-nt SRCs expression; (F/H) 24-nt SRCs expression. Parents have great different high expressed SRCs, and gained by their hybrid in B and C cross combinations, especially in 22-nt SRCs (Figure S4A/E). More SRCs tendency downregulated in higher fold changed level (Figure S4C/D/G/H). However, this phenomenon was not show in A cross combination.

**Figure S5.**
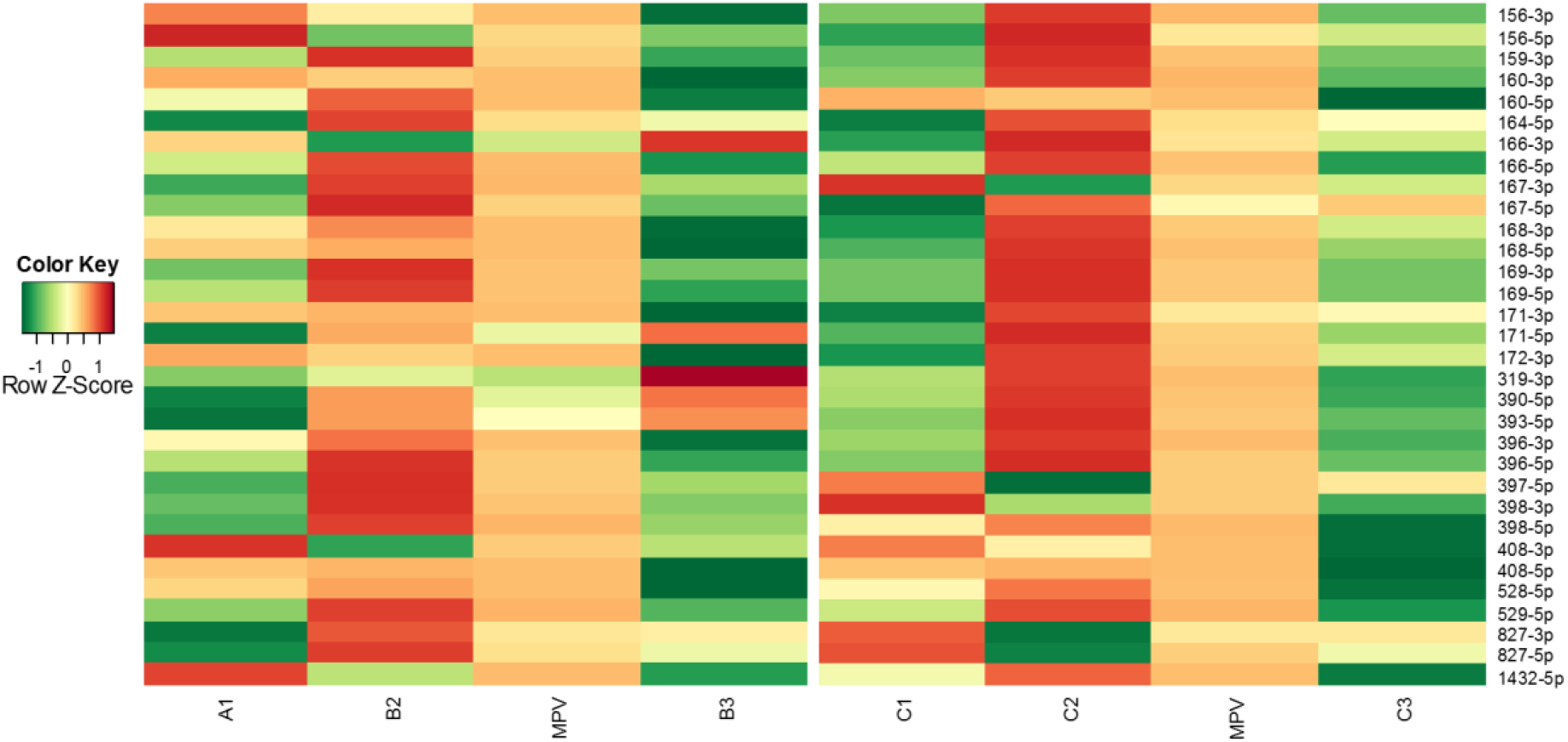
Known miRNA family’s expression in B and C cross combinations. Compare with middle parent value (MPV), most of the miRNA family downregulated in B and C cross combination.

**Figure S6.**
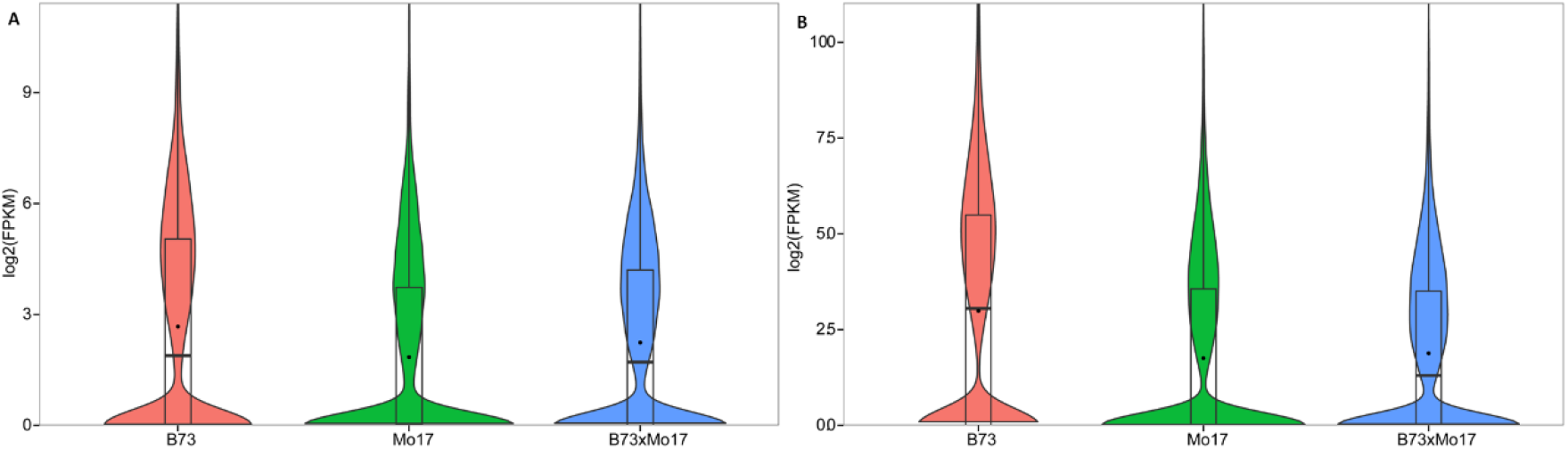
Expression of DNA transposon from 24-nt SRC regions (A) root (B) shoot

